# Folding and knotting of biotic and pre-biotic amino acid sequences through reverse evolution

**DOI:** 10.1101/2025.06.04.657801

**Authors:** João N. C. Especial, Patrícia F. N. Faísca

**Affiliations:** BioISI – Instituto de Biossistemas e Ciências Integrativas, Departamento de Física, Faculdade de Ciências, Universidade de Lisboa, 1749-016, Lisboa, Portugal

## Abstract

We develped a simple reverse evolution method to explore the protein folding transition and knotting process in globular proteins throughout evolution using as a proxy for evolutionary time the lenght of the amino acid alphabet. Three small proteins were considered. An unknotted one featuring a beta-sandwhich fold (FN3), a protein embedding a shallow trefoil knot (MJ0366), and a deeply knotted trefoil protein (YibK). Results from Monte Carlo simulations of a native-centric C_*α*_ model show that thermal stability increases througout evolution for all considered proteins suggesting it provided a selective pressure for the introduction of biosynthetic amino acids in the alphabet repertoire. Additionally, the thermodynamic cooperativity, the two-state nature of the folding transition, and the kinetic efficiency of folding (and knotting) displayed by the unknotted (and shallow knotted) protein have been roughly conserved throughout evolution, indicating that early alphabets, and, in particular the anscestral one composed of 10 amino acids, is competent to self-assemble relatively complex native structures. However, while the ancestral alphabet is clearly more efficient in collapsing the polypetpide chain to non-specific (i.e. non-native) knots, it significantly compromises the thermodynamic cooperativity and the folding ability of the deeply knotted protein YibK. The findings presented here collectively suggest that the alphabet of amino acids evolved to improve folding, but early alphabets might not have favoured the folding of native structures with deep knots. This could help to explain why deep knots in proteins are statistically uncommon.

## Introduction

Knotted proteins are proteins whose native structure embeds a physical knot (i.e., an open knot). The first knotted protein was reported in 1977,^1^ but it was only in 2000,^2^ following the development of computational methods to detect knotted topologies in open polymer chains, that these intricate molecules came into the spotlight. Nowadays, it is known that knotted proteins represent 1% of all protein entries available in the Protein Data Bank (PDB),^3^ being considered statistically rare when compared with random compact loops of the same size. ^4^

Knots in proteins are classified based on a topological invariant, which is the minimal number of crossings of a planar projection of the chain. The 3_1_ (or trefoil) knots, featuring three crossings, are the most ubiquitous, followed by 4_1_, and 5_2_ knots, with four and five crossings, respectively. Depending on its localization within the polypeptide chain, a protein knot classifies as deep if it is conserved after removing 20 residues from both termini;^5^ otherwise it is considered shallow. Deep trefoil knots are far more prevalent than knots with more crossings.^3^

The presence of knots in protein structures leads to increased folding complexity. Indeed, deeply knotted proteins often populate intermediate states (both in bulk and in single molecule studies) and exhibit multiple folding pathways. ^6^ Because of their complex folding mechanisms, the kinetics is often also complex, featuring distinct phases. Molecular simulations have shown that knotting proceeds through an ordered sequence of events where one of the protein termini threads a pre-formed twisted loop.^7^ This ordered process is a major source of backtracking, which slows down the kinetics or even impedes correct folding due to topological trapping.^8,9^

*In vivo*, the process of protein folding is often assisted by the so-called chaperonins; these are large, multi-subunit protein complexes that act by encapsulating unfolded or misfolded proteins within their central cavity, where the folding process occurs in an isolated and controlled manner, increasing folding effectiveness (i.e. the probability of a successful outcome).^10^ Molecular simulations demonstrated that protein encapsulation can effectively reduce the entropic cost of knotting, thereby lowering the folding free energy barrier.^11,12^ Furthermore, a seminal study based on an experimental method that probed protein folding directly upon synthesis on the ribosome showed that the bacterial chaperonin GroEL-GroES accelerates the folding of deeply knotted proteins YibK and YbeA by at least one order of magnitude, thus making their folding process more efficient.^13^

Knotted proteins exist in the three domains of life, which indicates their evolutionary significance and suggests that they may have evolved to play a specific biological role.^14^ Studies based on computer simulations suggested that knots can increase thermodynamic, kinetic, mechanical, or other form of native state stability, but experimental studies carried out so far have shown that all these forms of stability are within the same range for both knotted and unknotted proteins.^15^ Simulations and experiments agree that the knotted structure adopts a unique way of binding ligands in the SPOUT class of methyltransferases, indicating a role for the knot in catalysis, at least within this protein family.^16^ Recently, through a combination of simulations and experiments, it was proposed that the knotted topology in protein UCH-L1 contributes to the stabilization of the secondary structural elements. ^17^ However, determining the precise function of knots in proteins remains a puzzling problem, and one should not rule out the possibility that in general they serve no functional purpose at all.

The lack of a clear functional advantage associated with less efficient folding processes begs the question: Why are there knotted proteins at all? One explanation could be that knotting *in vivo* has always been assisted by chaperonins, and other cellular structures (e.g. the ribosome^18^) that provide spatial confinement. But then the question becomes: Why aren’t knotted proteins more frequent?

It is known that knotting frequency in polymeric systems can be modulated by physical properties (e.g., chain length and flexibility).^11,19,20^ Interestingly, it was found through extensive computer simulations of hetero patchy polymers that the alphabet size (i.e., the number of different types of monomers), and the monomer’s arrangement within the chain, can also significantly affect the knotting frequency.^21^ Smaller alphabets yield a higher abundance of knots compared to larger alphabets, and increasing the alphabet to 20 letters tends to suppress knots in these polymeric systems by allowing the formation of more complex local structures. These results are consistent with those obtained in the context of Monte Carlo simulations where it was proposed that a 20-letter amino acid alphabet may not be necessary to efficiently knot a lattice protein.^22^ Indeed, when the size of the amino acid alphabet was reduced by 30% the prevalence of knots increased by at least 25%. Altogether, these findings highlight the intricate interplay between polymer design, folding efficiency, and knotting behaviour.

Actual proteins are formed from a repertoire of the canonical set of 20 amino acids. However, there is broad consensus that only 10 of them (the less complex A, D, E, G, I, L, P, S, T, V) were present in prebiotic earth (i.e., in the absence of biology).^23^ Additionally, it is known that the 10 prebiotic amino acids were present at high enough levels to create functional polypeptides, ^23–25^ and that they optimally encode local backbone structures (i.e., secondary structural elements) and near-optimally encode tertiary contacts.^25^ The evolution of biosynthetic pathways, and the emergence of life, are associated with the appearance of the more complex, biotic (or modern) amino acids, which feature larger side chains allowing a broader range of proteins with diverse structures, functions, and properties.

Collectively, the results and observations outlined above suggest that a more profound understanding of knotted proteins must result from an evolutionary perspective on folding and knotting physics in relation to the length of the amino acid alphabet. Investigating this relationship within the framework of computer simulations is the aim of the present work.

## Model systems

To explore how the folding of knotted proteins developed through evolution we consider two proteins featuring a trefoil knot in their native structures. Protein MJ0366 from *Methanocaldococcus jannaschii* is the smallest knotted protein found to date. It has 92 residues, and the knotted core comprises residues 11 to 82 (Figure 1 A). Since both knot tails are short (10 residues), the knot is classified as shallow. The second protein is YibK, a 160 residue homodimeric a methyltransferase from *Haemophilus influenzae*, whose knotted core extends from residue 77 to residue 119. In this case the N-tail is 76 residues long and the C-tail 41 residues long (Figure 1B). The knot classifies as deep. Here, we investigate the monomer of YibK. We aditionally consider an unknottted protein, namely, a Fibronectin type III domain (FN3) from tenascin, which is a small protein domain with 90 residues, featuring a *β*-sandwich fold (Figure 1 C).

**Figure 1.**
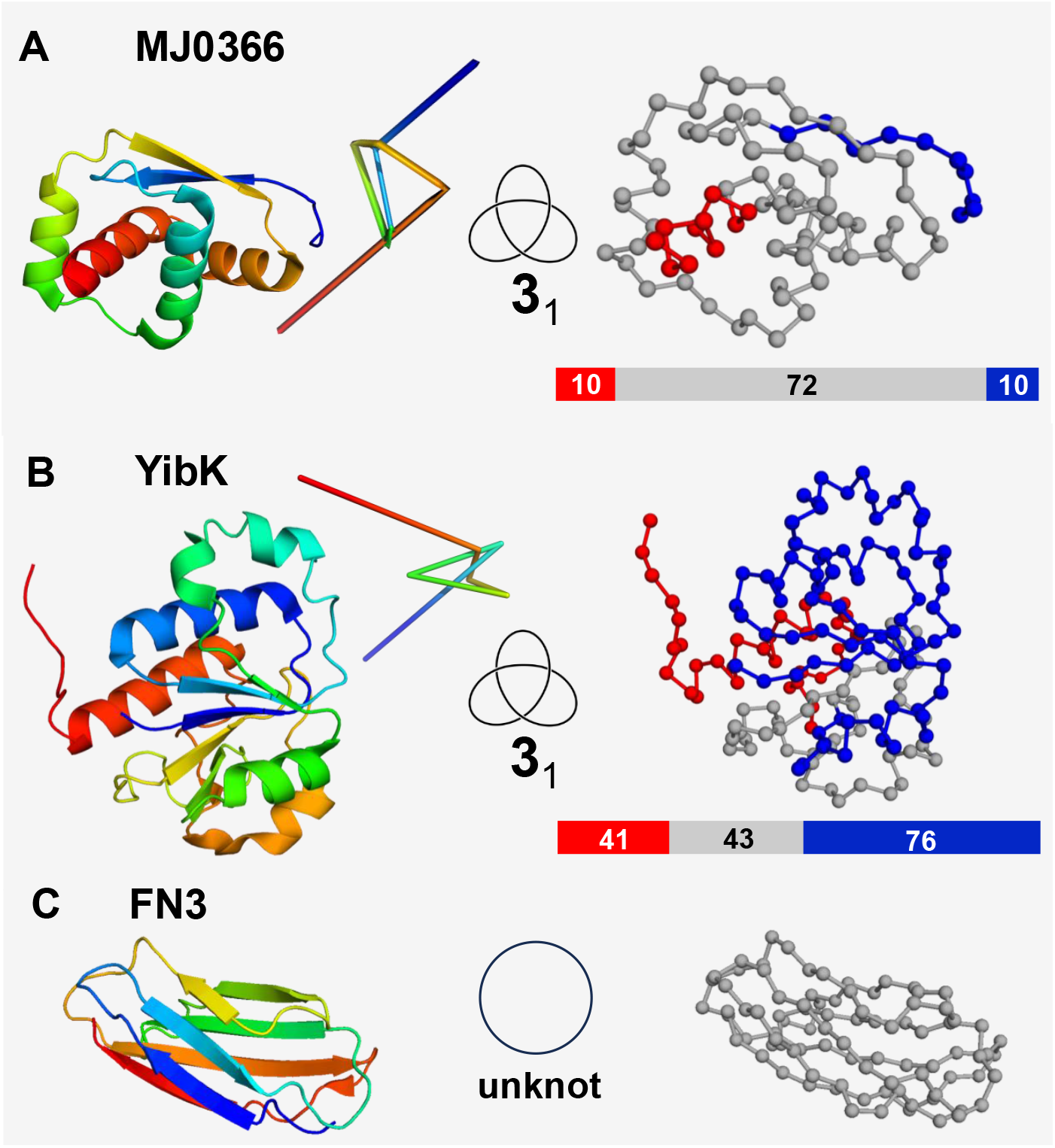
Model systems used in the present study. Cartoon representation (left) and bead and stick representation (right) of the native structure of proteins MJ0366 (PDB id: 2efv) (A), monomer of YibK (PDB id: 1j85) (B) and FN3 (PDB id:1ten) (C) with the knot tails coloured in red and blue. Each bead represents a C_*α*_ atom and rigid sticks represent pseudobonds connecting pairs of C_*α*_ atoms. The size (measured in number of beads) of the knot tails and knotted core is indicated. For the trefoil knotted proteins a reduced knot representation that results from applying the Koniaris-Muthukumar-Taylor (KMT) algorithm^2^ is also shown, as well as the knot localization within the polypeptide chain.

## Model and Methods

### The C_*α*_ Gōmodel

Proteins are represented by a simple C_*α*_ model (Figure 1). Accordingly, residues are reduced to hard spherical beads of uniform size, centered on the C_*α*_ atoms. Consecutive C_*α*_ atoms are connected by rigid sticks representing pseudobonds on the amide planes. We adopt a radius of 1.7 Å for the beads, which is the van der Waals radius of C_*α*_ atoms,^26^ and for the length of each stick we adopt the distance between the C_*α*_ atoms of the respective bonded residues in the protein’s native conformation, these being approximately 2.9 Å, for cis bonds, and 3.8 Å, for trans bonds. Two non-bonded residues are said to be in contact in the native conformation if the smallest distance between any two heavy atoms, one belonging to each residue, is 4.5 Å, this cut-off being chosen because it is slightly larger than twice the average van der Waals radius of heavy atoms in proteins.

To model protein energetics we consider the native-centric Gō potential.^27^ Accordingly, the total energy *E* of a conformation defined by bead coordinates 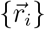 is given by

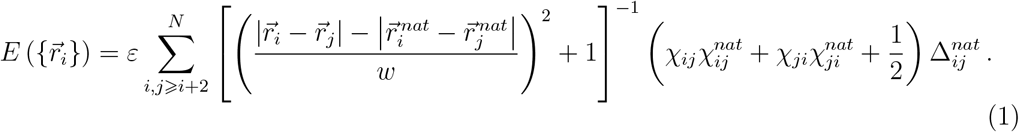

where *N* is the chain length measured in number of beads, 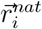 is the position vector of bead *i* in the native structure, 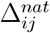 is 1 if the *i* − *j* contact is present in the native conformation and is 0 otherwise, *ε* is a uniform intramolecular energy parameter (taken as −1 in this study, in which energies and temperatures are shown in reduced units), *w* is the half-width of the inverse quadratic potential well, and the chirality of contact *i* − *j* in the conformation under consideration is

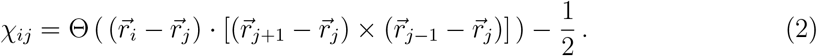

The chirality of the *i* − *j* contact in the native conformation is

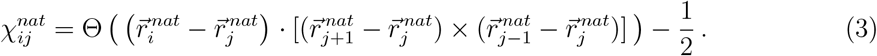

In equations (2) and (3), Θ is Heaviside’s unit step function, which takes the value 1 if its argument is greater than zero and the value 0 otherwise. The chirality factor in (1) favors the native conformation *vis a vis* its mirror conformation. A native contact is considered formed if the distance between the centers of the respective beads differs from the distance between their C_*α*_ atoms in the native conformation by less than the half-width of the potential wells, *w*.

### Replica-exchange Monte Carlo simulations

The conformational space of the C-alpha model is explored with Metropolis Monte Carlo (MC)^28^ by means of a move set that comprises crankshaft and pivot moves. All simulations start from an unfolded (and unknoted) conformation obtained from a simulation at high temperature, and folding progress is monitored by using the fraction of native contacts *Q*. In order to sample equilibirum distributions, we use Monte Carlo replica-exchange (MCRE).^29,30^ In this case the deployed move set does not preserve the linear topology of the chain (i.e., it allows the chain to cross itself) because the corresponding simulations require considerably less MC steps to equilibrate than those based on a move set that does not allow the chain to cross.^31^ The weighted histogram analysis method (WHAM)^32^ is used to analyze data from the MC-RE simulations and produce maximum likelihood estimates of the density of states from which expected values for thermodynamic properties are calculated as functions of temperature. An example is the heat capacity, *C*_*V*_, defined in reduced units as *C*_*V*_ = (*< E*^2^ *>* − *< E >*^2^)*/T* ^2^. The melting temperature, *T*_*m*_, is determined as the temperature at which the *C*_*V*_ peaks. We measure the cooperativity degree of the transition by the ratio of the full width at half maximum (*FWHM*) of the *C*_*V*_ peak to the melting temperature, and the half-width of the potential well, *w*, is adjusted to obtain a simulated *FWHM/T*_*m*_ ratio between 4 and 5%. This criterion has been successfully used in previous simulations employing a similar potential and sampling. ^11,12,33,34^ WHAM is additionally used to project the density of states along *Q* to obtain free energy and knotting probability profiles at a specific temperature. We define knotting probabiliby, *p*_*K*_, as the fraction of knotted conformations found in an ensemble of conformations (e.g., sharing the same *Q*).

### Fixed temperature Monte Carlo simulations

The number of Monte Carlo steps (mcs) does not correspond to physical time. Thus, to have a measure of folding efficieny, instead of considering the folding time (or folding rate) we decided to evaluate the foldicity, i.e., the number of unfolded-to-folded transitions that occur per million mcs (Mmcs) at fixed temperature *T < T*_*m*_ (i.e. in conditions that stabilize the native state). A model system that folds more efficiently will exhibit a higher number of such transitions at a certain temperature. For the purpose of evaluating folding efficiency the deployed move set must preserve the linear topology of the chain, i.e., it should not allow the chain to cross itself. As in the equilibrium simulations, a fixed temperature simulation also starts from a conformation that is unfolded (and unknoted). However, the simulation now stops when it reaches the native conformation. A conformation is considered folded if it is knotted, and its *Q* is larger than that at which the knotting probability is 0.99. Apart from measuring foldicity we also measure the number of knotting transitions per Mmcs to the first knotted conformation, and the number of knotting transitions per Mmcs to the last knotted conformation sampled prior to the native conformation is reached; we designate these by knoticity 1 and knoticity 2, respectively.

### Topological state

In both equilibrium and fixed temperature simulations the topological state (knotted or unknotted) of a sampled conformation is determined using the Koniaris-Muthukumar-Taylor (KMT) algorithm.^2^

## Results and discussion

### Reverse evolved sequences

To explore the physics of protein folding and knotting in a evolutionary context, we use a simplified model of reverse evolution to create primary structures made up of reduced alphabets. Specifically, the modern primary structure, which uses an alphabet of 20 amino acids, is gradually reverted through amino acid substitution to an hypothetical ancestral primary structure that can only use in its composition the 10 prebiotic amino acids (G, V, T, L, I, S, A, E, P, and D). Each intermediate sequence represents an evolutionary step and the lenght of the amino acid alphabet represents a stage of protein evolution serving as a proxy for evolutionary ‘time’. To perform amino acid substitutions we devised a simple criterium (which we designate by ‘revol’) that is based on a statistical reasoning that compares the relative abundance of the 20 naturally occurring amino acids in the biotic (B) and prebiotic (P) eras (Table 1).^35^

**Table 1:** Relative abundances of the 10 prebiotic amino acids (G, V, T, L, I, S, A, E, P, D) and their variation from the prebiotic era (P) to the biotic era (B) compared to the present relative abundances of biotic amino acids (R, K, N, Q, F, Y, H, M, W, C).

|  | **Prebiotic AAs** |  |  | **Biotic AAs** |  |
| --- | --- | --- | --- | --- | --- |
|  | relative abundance<br>P | relative abundance<br>B | variation<br>(P - B) | relative abundance |  |
| **D** | 0.05 | 0.05 | 0.00 | 0.00 | **C** |
| **P** | 0.07 | 0.05 | 0.02 | 0.01 | **W** |
| **E** | 0.08 | 0.06 | 0.02 | 0.02 | **M** |
| **A** | 0.12 | 0.09 | 0.03 | 0.02 | **H** |
| **S** | 0.10 | 0.07 | 0.03 | 0.03 | **Y** |
| **I** | 0.10 | 0.06 | 0.04 | 0.04 | **F** |
| **L** | 0.14 | 0.10 | 0.04 | 0.04 | **Q** |
| **T** | 0.10 | 0.06 | 0.04 | 0.04 | **N** |
| **V** | 0.12 | 0.07 | 0.05 | 0.05 | **K** |
| **G** | 0.12 | 0.07 | 0.05 | 0.06 | **R** |

Specifically, we consider the relative abundance of the biotic amino acids, and the difference between the (estimated) relative abundance of the prebiotic amino acids in the prebiotic and biotic eras, i.e., (P-B). If we order prebiotic amino acids by increasing values of the (P-B) variation, and biotic amino acids by increasing values of their present relative abundance, we find that on the same line these two values are always quite close to each other (Table 1). This suggests a highly simplified hypothesis for protein evolution: that each biotic amino acid (on the rightmost column) replaced only instances of the prebiotic amino acid on the same line (on the leftmost column) in a stepwise (i.e. gradual) manner. Such a mechanism of amino acid replacement would be compatible with the codon capture mechanism for the evolution of the genetic code.^36^ Codon capture occurs when a tRNA that was previously responsible for carrying a specific amino acid gains the ability to carry a different amino acid. It underlies a gradual expansion of the genetic code by allowing new amino acids to ‘capture’ codons that were previously used by other amino acids. The gradual introduction of amino acids into the genetic code suggests that new amino acids were added over time, rather than all at once. If we further assume that larger coarser changes occured earlier in evolution than finer incremental ones, larger (P-B) variations should have occurred earlier in the evolution process and evolution would have progressed from bottom to top of Table 1. This hypothesis thus suggests that the sequence in which biotic amino acids may have been introduced could have been: R-K-N-Q-F-Y-H-M-W-C. Interestingly, this sequence is quite similar to that proposed by Liu et al.,^37^ K-R-N-F-Q-Y-M-H-W-C, based on a statistical analysis of amino acid usage across species. On the basis of this hypothesis, we can ‘reverse evolve’ (hence the designation ‘revol’ for this procedure) a current primary sequence in a stepwise manner by (from top to bottom of Table 1) first reducing its alphabet size from 20 to 19 by replacing all cysteines (C) by aspartic acids (D), then from 19 to 18 by replacing all tryptophans (W) by prolines (P), and so forth until, when all arginines (R) are replaced by glycines (G), the alphabet size becomes 10 (Figure 2A).

**Figure 2.**
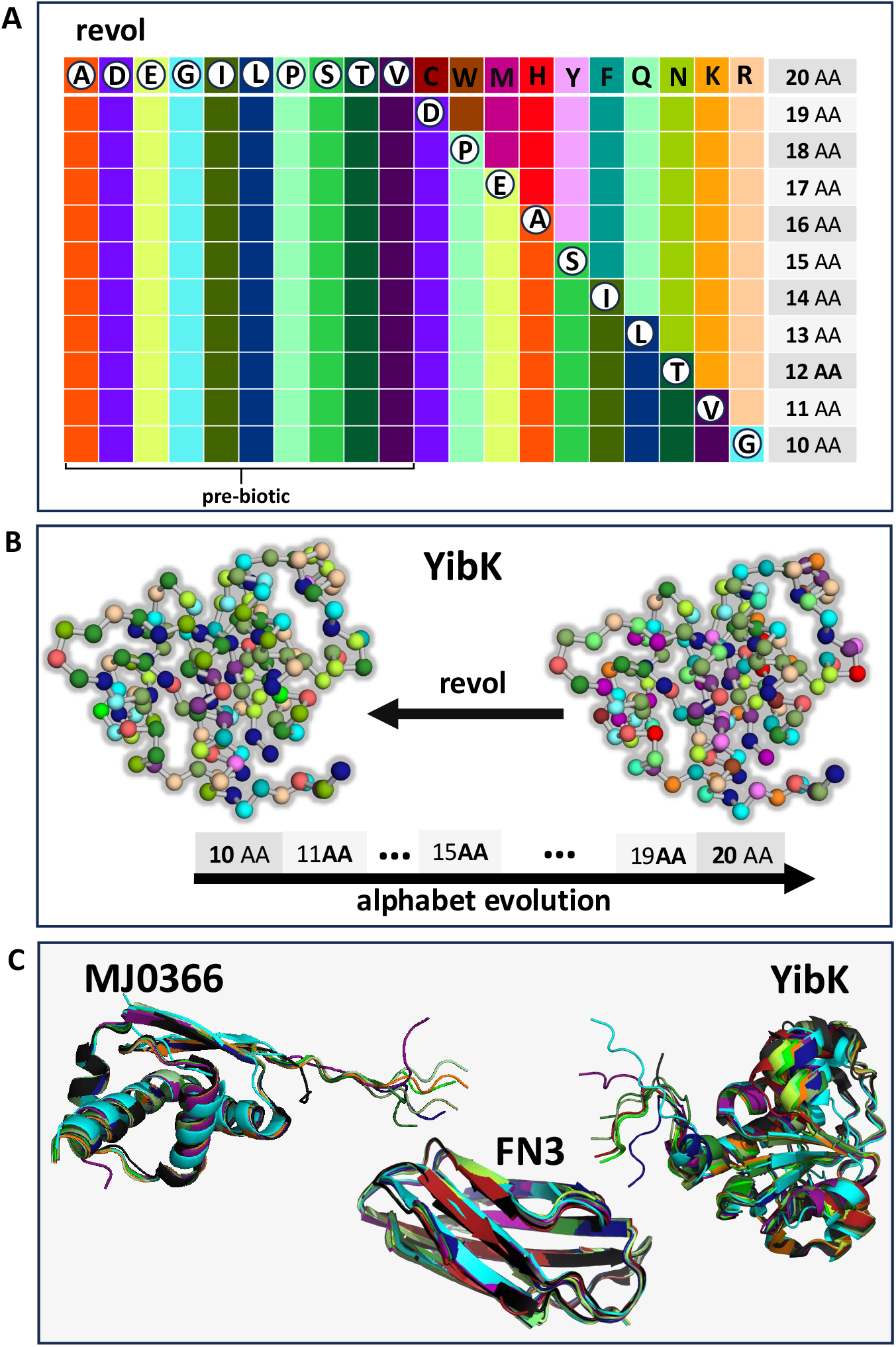
Reverse evolution. (A) Order according to which the amino acid substitutions are performed based on the revol criterion. (B) Modern (right) and ancestral (left) YibK. (C) Superposition of the 9 (MJ0366), 10 (YibK) and 8 (FN3) native structures generated by AF on the corresponding PDB structures to highlight the structural similarity quantified by the RMSD.

The primary sequence of MJ0366 lacks tryptophan (W) residues, and that of FN3 lacks both cysteine (C) and histidine (H). Therefore, once the single cysteine (C) of MJ0366 is replaced by an aspartic acid (D), the alphabet size drops from 20 to 18 in the first step of reverse evolution. In the case of FN3, the process of reverse evolution skips the first and forth steps. Since the number of amino acids decreases from 20 to 10 when going from the biotic to the prebiotic alphabet, the application of each criterion to the the modern primary sequence of MJ0366, YibK and FN3 generates 9, 10 and 8 precursor primary sequences, respectively, each one representing an evolutionary stage. The sequence resulting from the first amino acid substitution performed in the modern sequence corresponds to the precursor prior to the last evolutionary step, and the one resulting from 10 amino acid substitutions is the ancestral sequence (Figure 2B).

### Reverse evolved native structures

To find the native structures corresponding to the precursor primary sequences generated by the application of revol (featuring 1, 2, 3, 4, …, 10 substitutions) we tested three structure predictions tools: AlphaFold2,^38^ the version of AlphaFold2 available in ColabFold, ^39^ and ESMFold.^40^ A major difference between AlphaFold2 and ESMFold is that the former uses multiple sequence alignments (MSAs) and templates of similar proteins to achieve optimal performance, while the latter generates structure prediction using only one sequence as input by leveraging the internal representations of the language model. For each generated primary sequence the output of AlphaFold2 comprises 5 native structures, with the first one being the best prediction (i.e., the one with the highest confidence).

The native structures predicted with AlphaFold2 for all generated primary structures (which have alphabet sizes ranging from 10 to 20 amino acids) all embedded the same trefoil knot. However, both the ColabFold (in the structure to which it assigned most confidence) and ESMFold, incorrectly predicted the topological state of the (modern) primary sequence of protein MJ0366. In view of this outcome, we adopted AlphaFold2 as the structure prediction tool for this work and refer to it simply as AlphaFold (AF). To further compare the native structures predicted with AF we used PyMol^41^ to calculate the root-mean-square-deviation (RMSD) at the level of C-alpha atoms to the PDB native structure. In the case of MJ0366, the RMSD ranges from 0.47 Å (13 letter alphabet) to 1.09 Å (10 letter alphabet), from 0.79 Å (18 letter alphabet) to 3.15 Å (10 letter alphabet) in the case of YibK (Figure 2C), and from 0.33 Å (20 letter alphabet) to 1.055 Å (13 letter alphabet) in the case of FN3. It is interesting to note that the native structures predicted by AF for the modern alphabet have consistently more native contacts than the natural ones.

### Number of native contacts

We started by investigating the relation between alphabet size and the number of native interactions. The (arithmetic) mean number of atoms in the side chains of prebiotic amino acids is 7.9, whereas in the case of biotic aminoacids this number incrases to 12.5. Since the construction of the native contact matrix for the Gō potential is based on a full atomistic protein representation, early alphabets, which have overall amino acids with less atoms, are expected to give rise to a native contact matrix with fewer contacts. We find that the number of native contacts increases linearly with the size of the amino acid alphabet for YibK, and the dependence is moderately linear for the other model proteins (Figure 3). The modern native structure of YibK contains 14% more native contacts than YibK’s ancestral native structure. In the case of MJ0366 and FN3, the increase in the number of native contacts is 8% and 9%, respectively.

**Figure 3.**
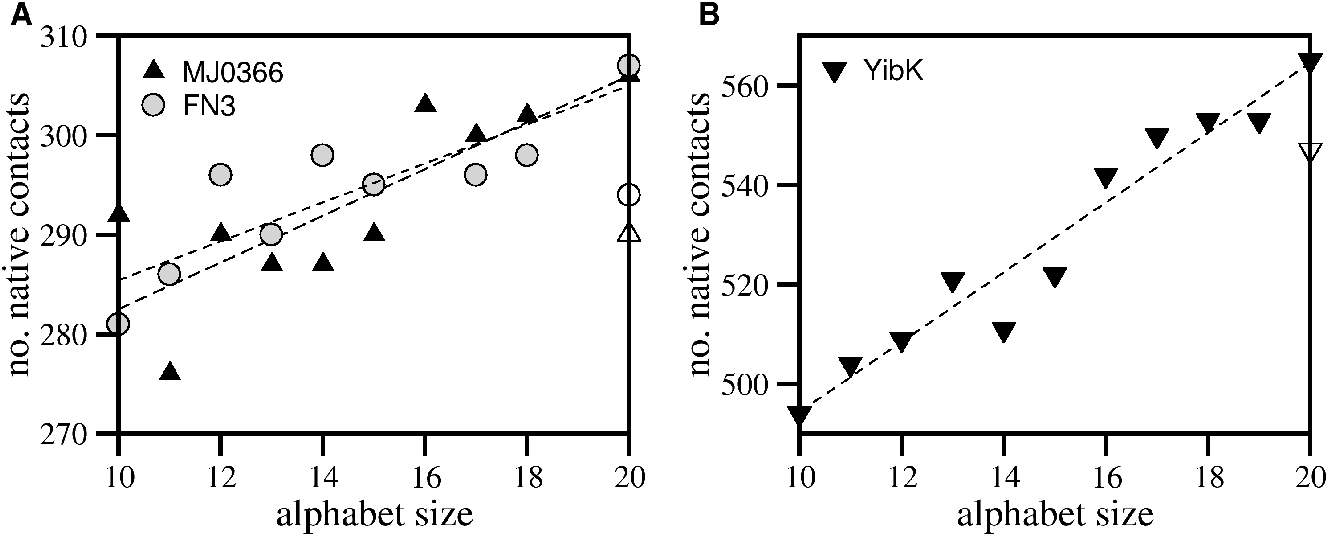
Number of native contacts and alphabet size. The number of native contacts increases linearly with the size of the amino acid alphabet for protein YibK (*R* = 0.97, *t* = 12.26, null hypothesis *p* = 6.44 *×* 10^−07^). For MJ0366 (*R* = 0.82, *t* = 3.907, null hypothesis *p* = 0.0045) and FN3 the correlation is moderate (*R* = 0.86, *t* = 4.618, null hypothesis *p* = 0.0024). The number of native contacts corresponding to the PDB structure is colored white.

### Thermal stability

The melting temperature (*T*_*m*_) is a measure of protein thermal stability and corresponds to the temperature at which the *C*_*V*_ peaks. In the context of the Gō potential, the energy of the native state is direclty proportional to the number of native contacts, and the latter are all stabilizing. Consequently, the native structure should become energetically more stable as the size of the alphabet increases, and, therefore, the thermal stability should increase with alphabet size. Figure 4 (A-C) reports the dependence of the *C*_*V*_ on *T*. Interestingly, the heat capacity curve corresponding to the ancestral YibK protein exhibits two peaks (a smaller one at *T* = 0.77, and a larger one at *T* = 0.88, which we identify as the *T*_*m*_), which is an indication that it does not denature via a two-state transition, populating detectable thermal intermediate states (as shown ahead the one at lower temperature is knotted, while the other is unknotted). Additionally, we find that the relation between *T*_*m*_ and alphabet size is roughly linear for FN3 and MJ0366 (Figure 4 D, E), but distinctively sigmoidal in the case of YibK, with the transition midpoint observed between alphabet size 15 and 16 (Figure 4 F).

**Figure 4.**
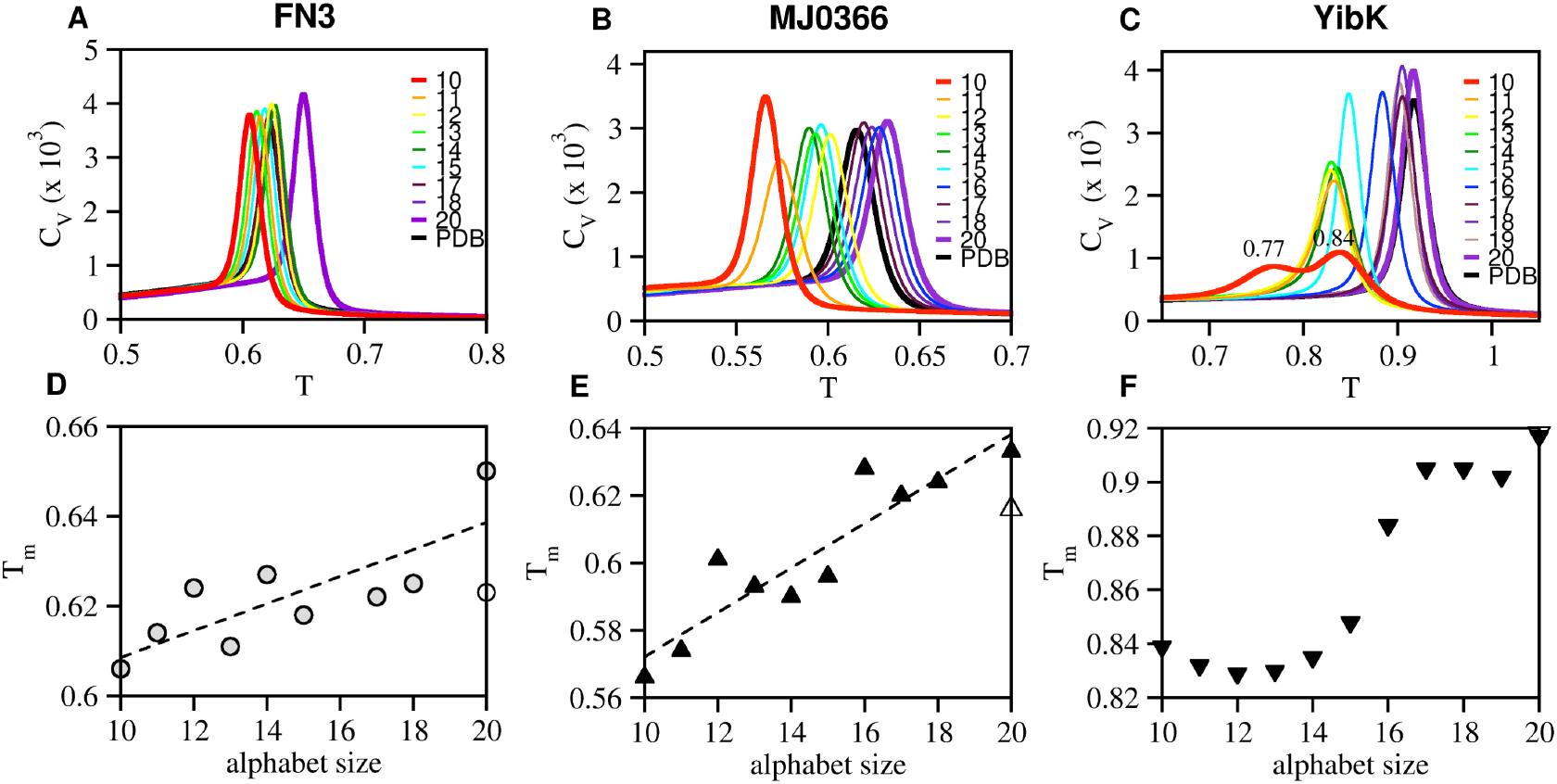
Thermal stability. Dependence of the heat capacity, *C*_*V*_, on the temperature, *T* for FN3 (A), MJ0366 (B) and YibK (C). In the case of YibK we indicate the temperatures corresponding to the two peaks of the *C*_*V*_, with the largest peak being identified with *T*_*m*_. Dependence of the melting temperature *T*_*m*_ on the size of the alphabet (D-F). The *T*_*m*_ corresponding to the PDB structure is colored white.

### Thermodynamic cooperativity

To quantify the degree of thermodynamic cooperativity (i.e., thermodynamic two-state-like behavior of the folding transition) a dimesionless quantity termed van’t Hoff ratio, *k*_2_ = Δ*H*_*vH*_*/*Δ*H*_*cal*_, is often used both in simulations and experiments. In the definition of 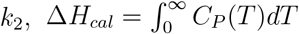 is the calorimetric enthalpy, and 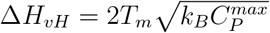 is the van’t Hoff enthalppy. The latter corresponds to twice the maximum standard deviation of the enthalpy distribution at the temperature at which the *C*_*V*_ is maximum. A *k*_2_ close to unity is typically taken as an indication that the folding transition is thermodynamically cooperative, taking place in an apparent two-state manner (i.e., with a negligible population of intermediate states at *T*_*m*_).^42^ We note that in the context of the present model the values of *H* and *C*_*P*_ are essntially identical to those of *E* and *C*_*V*_ obtained in the scope of the adopted simulational protocol, and we used them to evaluate *k*_2_. The results from simulations (Figure 5) indicate that for MJ0366 and FN3 a *k*_2_ *≈* 1 is conserved across evolution. Not surprisingly, in the case YibK, the ancestral protein exhibits a rather small value of *k*_2_ = 0.33, which is consistent with the double peaked *C*_*V*_ curve. Moreover, for YibK, the van’t Hoff ratio only attains values close to unity when there are at least 15 different amino acids available to make up the protein’s primary sequence (Figure 5).

**Figure 5.**
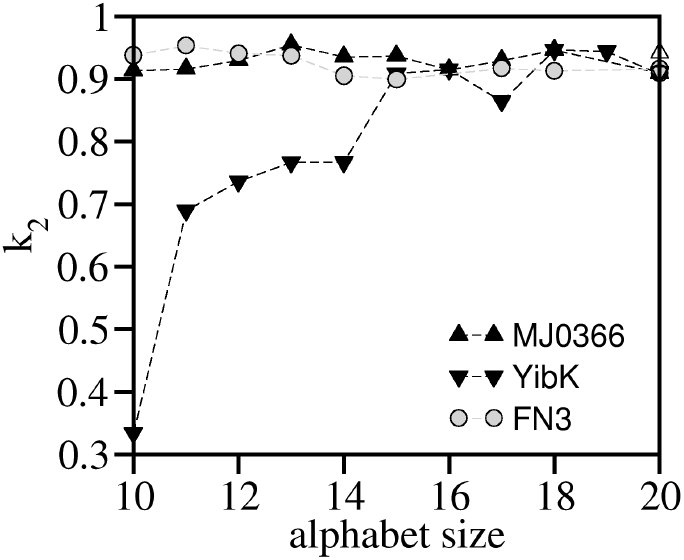
Thermodynamic cooperativity. Dependence of the van’t Hoff ratio, *k*_2_, on the alphabet size for the three studied proteins. The van’t Hoff ratio corresponding to the PDB structure is colored white and for the three model systems coincides with that of the model protein predicted by AF.

### Free energy

For each considered alphabet, we projected the free energy on the reaction coordinate *Q* at the corresponding *T*_*m*_. Aditionally, in the case of YibK we also report the dependence at *T* = 0.77, which is the temperature at which the heat capacity firstly peaks. The free energy profiles obtaind for FN3 and MJ0366 are consistent with a two-state folding transition, which is conserved throughout evolution (Figure 6 (A-C). However, in the case of YibK, the transition is clearly two-state only when the amino acid alphabet contains at least 14 different letters, despite the free energy barrier decreasingly significantly from size 15 to 14 (Figure 6 (C)). Indeed, smaller alphabets render smaller free energy barriers, with the transition becomming nearly downhill for the ancestral protein. It is interesting to note that protein does not fold to a knotted state at *T*_*m*_, and that even at the lowest temperature the probability to be knotted does not exceed 0.7 (Figure 6 (C, E), suggesting that native contacts that are critical to fold and knot to the native state were lost when moving from alphabet size 14 to 10. To explore this scenario we computed the contact maps of the native structures corresponding to alphabet sizes 20, 14, and 10, as well as those of the corresponding differentials (i.e., 20-14, 20-10 and 14-10). We find that the contact map is roughly conserved for alphabet size 14, but it changes significantly for alphabet size 10, especially between amino acids 80 and 150 (data not shown). Additionally, in the ancestral native structure there are 18 residues located in beta strands, against 30 residues found in the beta strands of the modern native structure (https://webclu.bio.wzw.tum.de/stride/). ^43^ This translates into loss of secondary structural content for the ancestral protein, and, since beta strands are mainly stabilized by long-range interactions, it also results into a loss of the latter. The decrease in secondary structural content, and in the number of non-local interactions can both contribute to a less cooperative folding transition. Indeed it has been suggested, based on molecular simulations, that folding cooperativity results from the interplay between local and non-local interactions,^42^ and from the interplay between secondary and tertiary structural content.^44^

**Figure 6.**
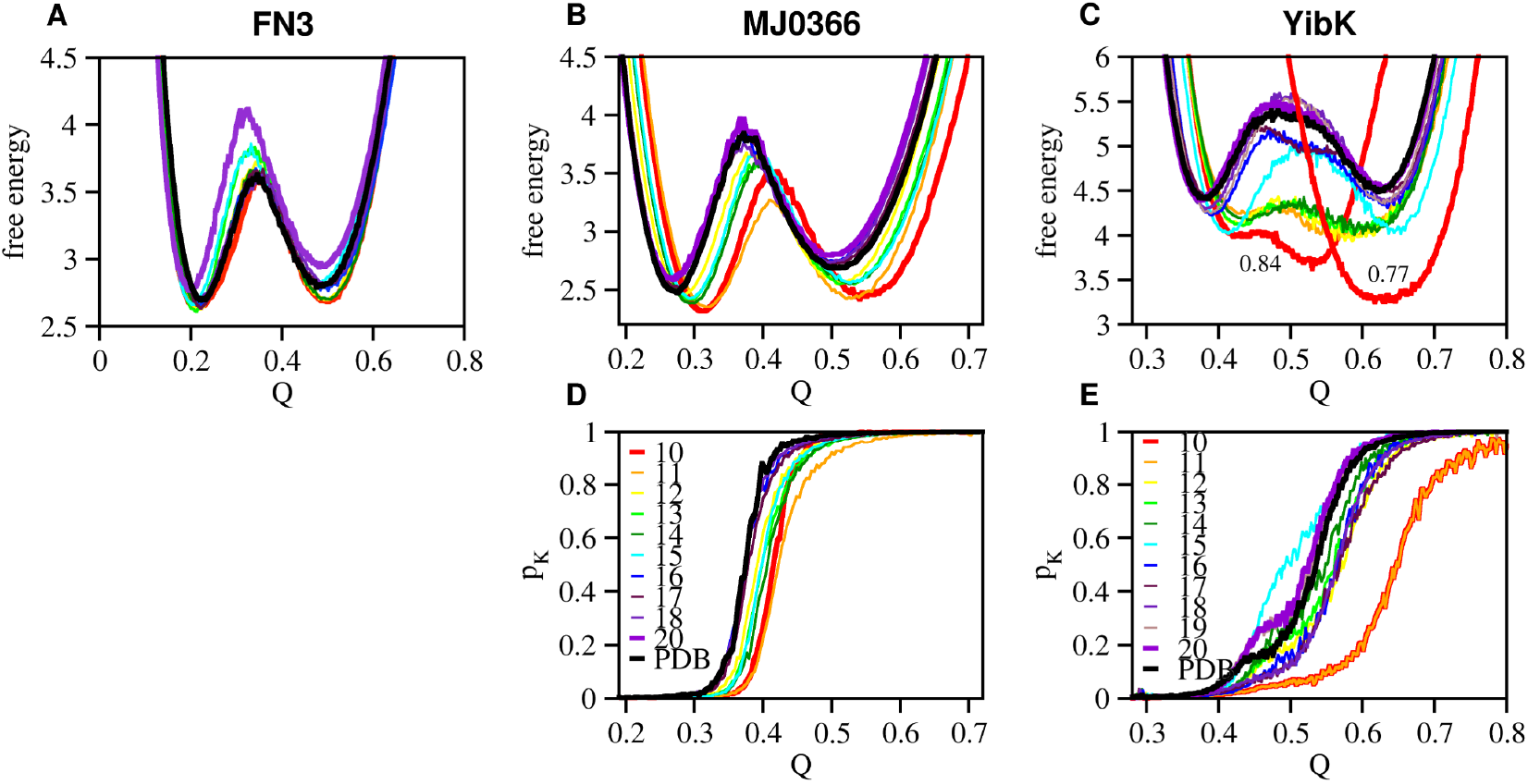
Free energy profiles and knotting probability profiles. Free energy projected on the reaction coordinate *Q* (A-C), and knotting probability projected on the reaction coordinate *Q* (D-E). In the case of YibK we also report the free energy profile corresponding to the first *C*_*V*_ peak (*T* = 0.77).

### Knotting probability

The projection of the knotting probability, *p*_*K*_, on *Q* (Figure 6 D, E) indicates that as the alphabet size decreases knotting tends to require a more structurally consolidated, and consequently, more energetically stable conformation, This effect is particularly striking in the case of the ancestral YibK (Figure 6 E), with the midpoint of the knotting transition inceasing 33% against the modern YibK.

### Folding efficiency

Here we investigate the efficiency of the folding process (measured as the number of recorded transitions to the native structure per Mmcs) at fixed temperature throughout evolution. In particular, for alphabet sizes 10, 12, 15, 17 and 20 we conduted a total of 20M fixed temperature transition simulations (all considered temperatures being below the *T*_*m*_ of the corresponding system). The results from simulations reveal a somewhat canonical behavior (which is conserved for alphabet sizes 12, 15, 17 and 20) for all considered model systems where there is one optimal temperature at which the folding efficiency is highest^45–47^ (Figure 7 A, B). On the other hand, in the case of YibK, this behavior breaks down and a rather distintct one appears; in this case the folding efficiency has one pronounced local minimum, and folding is observed within a rather shorter temperature range (Figure 7 C).

**Figure 7.**
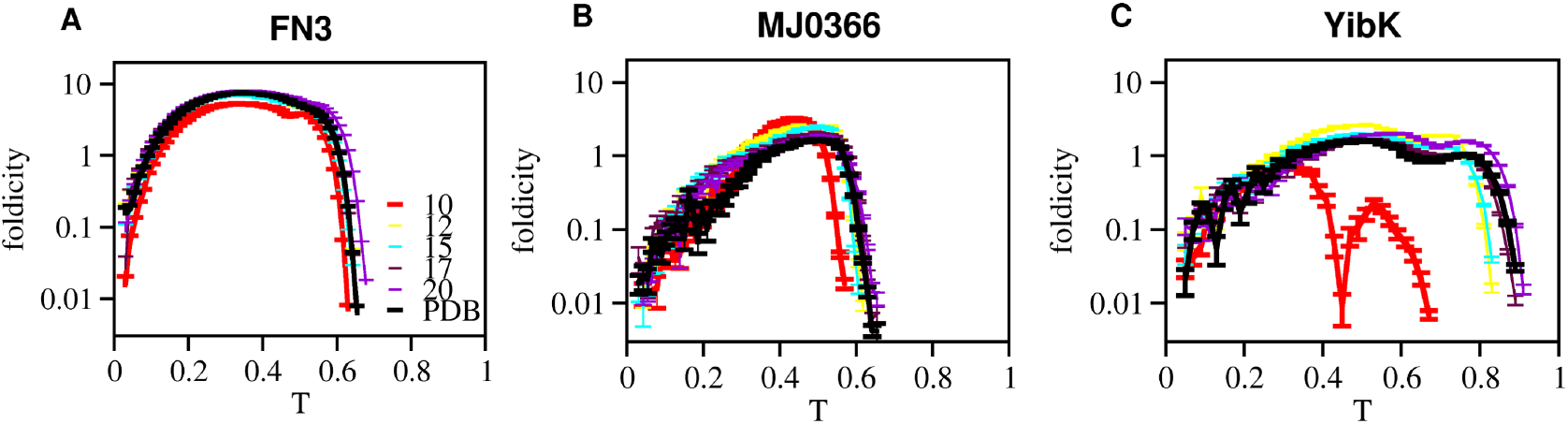
Folding efficiency. Dependence of (the logarithm of) foldicity (measured as the number of folding transition per Mmcs) for alphabet sizes 10, 12, 15, 17 and 20 for proteins FN3 (A), MJ0366 (B) and YibK (C).

### Knotting efficiency

Finally, we investigate the efficiency of the knotting process (measured as the number of knotting transitions per Mmcs recorded during folding to the native structure) at fixed temperature throughout evolution (Figure 8). The dependence of knotting efficiency measured by knoticity 1 (which concerns the first knotting event), shows a qualitatively similar dependence on themperature for all considered model systems, with the ancestral sequence of YibK knoting up to five times more efficiently than the other sequences, especially the modern one (Figure 8 A, B). Interestingly, the knotting efficiency mesured by knoticity 2 (Figure 8 C, D), which concerns the last knotting event before reaching native state, shows exactly the same dependence on temperature as the foldicity. This is a clear indication that in the considered model systems folding is limited by the formation of the native knot, which is, therefore, the limiting event.

**Figure 8.**
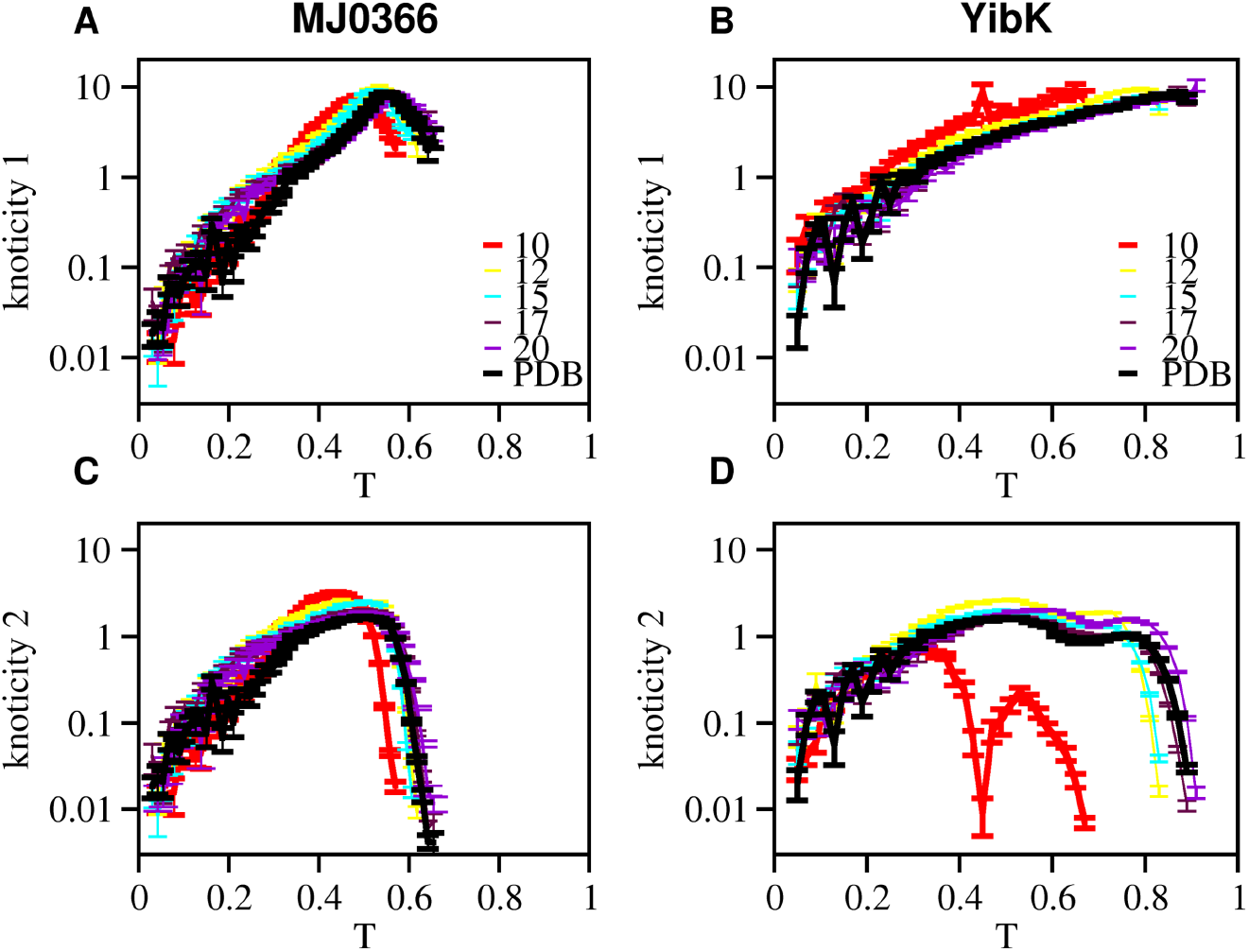
Knoting efficiency. Dependence of (the logarithm of) knoticity 1 (measured as the number of transitions to the first knot per Mmcs) and knoticity 2 (measured as the number of transitions to the last knot per MMC steps) on temperature for alphabet sizes 10, 12, 15, 17 and 20 for proteins MJ0366 (A, C) and YibK (B, D).

## Conclusions

This work studied the physics of protein folding and knotting in the context of a simple C_*α*_ native-centric Gō model throughout evolution. The adopted mechanism of protein reverse evolution is also simple, being based on a criterion that considers the relative adundance of amino acids in the biotic and pre-biotic eras. Starting from the moderm primary structure the application of this criterion generates a set primary sequences representing different evolutionary stages. While the latter actually represents an oversiplification that does not take into account other mechanims that contributed to the evolutoinary history of proteins (e.g., specific environmental factors, mutations, genetic drift etc.), it contributes to illuminate how amino acid availability may have influenced the development of folding physics and knotting behavior. The simplicitiy of the adopted Gō model, also implies that the the major feature of the amino acids influencing protein behavior is the size of the side chains, which directly affects the native contact map, and, consequently, protein energetics.

Three small proteins were investigated. An unknotted protein (FN3), a protein featuring a shallow knot (MJ0366), and another one featuring a deep knot (YibK). Native structures generated by AlphaFold for the biotic and pre-biotic primary structures were all found to be knotted suggesting that that even the earliest alphabet sizes are sufficient to encode knotted folds. This is consistent with the fact that knotted proteins are found in species from the most primitive archea superkingdom.

We found that all alphabets, including the ancestral one made up of 10 aminoacids, are able to drive a cooperative folding transition for both the unknotted and shallow knotted proteins, thus fullfiling the requirement of foldability (or the ability to fold into a specific conformation). This result is in line with a study based on the analysis of peptide folding, which concluded that the early alphabet composed of 10 amino acids was remarkably adaptive at supporting (cooperative) folding of the earliest proteins. ^48^ This observation is, however, not applicable to the ancestral deeply knotted protein YibK studied here that fails to exhibit a cooperative folding transition, collapsing downhill to a minimim free energy state that is not necessarily knotted. In other words, the ancestral pre-biotic alphabet composed of 10 amino acids, fails to fold a deeply knotted protein like YibK, at least in the context of the presently adopted C_*α*_ Gō model.

In all other model systems that fold cooperatively, we observe differences in the equilibirum properties of the folding transition. One such property, thermal stability, increases throughout evolution, although differently for the three considered proteins. The increase in thermal stability is likely the result of incorporating in the amino acid alphabet the larger side chains of the biotic amino acids. This led to an increase in the number of native interactions and, consequently, of the energetic stability of the native structure. The increase in thermal stability as a result of increasing the size of the amino acid alphabet may be one of the reasons why the alphabet actually evolved to incorporate biosynthetic amino acids.

We also found that other thermodynamic properties (thermodynamic cooperativity and free energy profiles) are conserved for the unknotted protein and protein with the shallow knot, being strongly dependent on the amino acid alphabet for the larger, deeply knotted protein. In particular, in this case, the projection of the free energy on the fraction of native contacts (at *T ⩽ T*_*m*_) is consistent with a downhill process to a free energy minimum which is not necessarily knotted.

The folding efficiency (measured by the number of folding transitions observed per million Monte Carlo steps) has a canonical dependence on temperature, with maximum folding efficiency for some specific temperature (below *T*_*m*_), for both the unknotted and shallow knotted protein. Additionally, this behaviour is conserved throughout evolution. However, in the case of YibK, the ancestral alphabet leads to a non-canonical folding dependence on temperature, and considerably lower folding efficiency. However, this alphabet size is actually more efficient than the modern one in knotting the protein to a non-specific conformation. It is thus possible that deeply knotted proteins require a larger amino acid repertoire to fold. This could, at least partly account for their statistical rarity.

## Author contributions

PFNF designed the research. JNCE developed the code, performed all the calculations and conceived the criteria used for reverse evolution. JNCE and PFNF analyzed the data. PFNF prepared the figures. PFNF and JNCE wrote the paper.

## Acknowledgement

Work supported by UID/00100, BioISI (DOI: 10.54499/UIDB/04046/2020) Centre grant from FCT, Portugal (to BioISI). JNCE acknowledges financial support from FCT, Portugal, through PhD grant SFRH/BD/144345/2019. A part of this work was performed on the computational resources of INCD (http://www.incd.pt) funded by FCT and UE under project LISBOA-01-0145-FEDER-022153. Access was granted by FCT through project 2022.26279.CPCA.A0.

